# HiCUP-Plus: a fast open-source pipeline for accurately processing large scale Hi-C sequence data

**DOI:** 10.1101/2022.05.18.492393

**Authors:** S. Thomas Kelly, Satoshi Yuhara

## Abstract

Hi-C is an unbiased genome-wide assay to study 3D chromosome conformation and gene-regulation. The HiCUP pipeline is an open-source tool to process Hi-C from massively parallel sequencing while accounting for biases specific to the restriction enzyme digests used. It is an excellent solution tailored to analyse this technique, however the latest aligner supported by the current release is Bowtie2. To improve the computational performance and mapping accuracy when using the HiCUP pipeline, we have modified it to optionally call the HiSAT2 and Dragen aligners. This allows using the HiCUP pipeline with 3^rd^ party aligners, including the commercially-licensed high performance Dragen aligner. The HiCUP+ pipeline is modified extensively to be compatible with Dragen outputs while ensuring that the same results as the original pipeline can be reproduced with the Bowtie or Bowtie2 aligners. Using the highly accurate HiSAT2 or Dragen aligners produces larger outputs with a higher proportion of uniquely mapped read pairs. It is therefore feasible to leverage the reduced compute-time of Dragen to reduce compute costs and turnaround-time without compromising quality of results. The HiCUP pipeline and Dragen both compute rich summary information.

## Background

Chromosome structure and conformation is highly spatially-organised and dynamic, playing a key role in gene regulation and DNA replication (Fraser et al., 2015). Understanding the 3-dimensional structure of chromosomes and how they interact is a question fundamental to basic biology and essential to applications in biotechnology, pharmaceuticals, and diagnostics. The “Hi-C” technique extends the chromatin conformation capture (3C) approachusing massively parallel genomic sequencing (Leibermann-Aiden, 2009) to identify pairs of sequences from genomic regions in close spatial proximity using ligation and restriction enzyme digests. Hi-C provides an unbiased genome-wide assay to map spatial contacts between chromosomal regions without prior knowledge of chromosomal organisation or putative interactions. The technique is quantitative, enabling the analysis of the frequency of interactions in the genome (Cameron et al., 2020). Hi-C is a ligation-based technique which generates pairs of sequences that cannot be expected to physically map to adjacent genomic loci. As such bioinformatics processing of the Hi-C data is challenging, requiring an unconventional approach of mapping each read to the genome independently (Wingett et al., 2015).

Due to its popularity in the field of regulatory genomics, Hi-C has been made commercially available (Arima, 2022; Dovetail Genomics, 2021; Qiagen, 2022). Large-scale functional genomics projects such as FANTOM6 (Ramilowski et al., 2020; Agrawal et al., 2021) have applied Hi-C to map cell type-specific domains. While originally developed with “6-cutter” restriction-enzymes, it is now recommended to use “4-cutter” restriction enzymes or DNase for higher resolution (Belaghzal et al., 2018; Cameron et al., 2020; Ma et al., 2018; Ramani et al., 2016). The technique has also been developed into single-cell Hi-C (Nagano et al., 2013) and used to demonstrate that localised regulatory structures are maintained between individual cells.

The Hi-C User Pipeline (HiCUP) is a bioinformatics pipeline specifically designed for processing Hi-C data (Wingett et al., 2015). HiCUP (Wingett et al., 2021) has been released open-source on GitHub (https://github.com/StevenWingett/HiCUP). The pipeline enables mapping Hi-C data to a reference genome, while removing technical artefacts such as removing adapter sequences, and correcting for biases specific to the cut sites of the restriction enzymes used. Hi-C analysis requires that each read is independently mapped to a unique locus in the genome with high accuracy. HiCUP is implemented in Perl and R having minimal dependencies, requiring only a Bowtie aligner, samtools, and some R packages to be installed. This allows HiCUP to be easily installed on a Linux server and ran in multiple threads with low memory requirements. While the pipeline supports either Bowtie (Langmead et al., 2009) or Bowtie2 (Langmead and Salzberg, 2012), the main drawback of the HiCUP pipeline is its reliance on these aligners, with poorer computational performance than numerous alternatives now available (Kim et al., 2015; Kim et al., 2019; Illumina, 2021).

## Development of HiCUP+

To address this, we first tested a customised version of HiCUP with the Hierarchical indexing for Spliced Alignment of Transcripts (HiSAT2) aligner (Kim et al,. 2019). We have modified the HiCUP source code to document additional aligners and allow configuration of new aligners and then updated the mapping step to test calling a new aligner. HiSAT2 was selected as a modern open-source algorithm using a novel indexing approach that is fast, sensitive, and widely available. It is one of the fastest and most accurate aligners currently available (Musich et al., 2021). HiSAT2 is implemented in Perl and supports stream inputs and outputs, as the currently supported Bowtie and Bowtie2 aligners. We were therefore able to show that custom aligners can be added to the HiCUP pipeline.

Next, we heavily modified the HiCUP pipeline to support the proprietary licensed Dynamic Read Analysis for GENomics (Dragen™) pipeline (Illumina Inc., 2021). Dragen uses hardware-acceleration on field-programmable gate array (FPGA) technology and optimised code implementations to drastically improve performance of alignment and various built-in quality-checking or summary statistics features. It is appealing to use Dragen for Hi-C analysis due to the clear benefits of improved computational performance and mapping accuracy compared to Bowtie2. However, Dragen is implemented in C++ and does not allow stream inputs. Therefore, it was necessary to extensively modify the HiCUP pipeline to run Dragen via a system call and then pass the results in Perl to downstream filtering and summary steps.

We have successfully implemented a modified HiCUP+ (HiCUP-Plus) pipeline, which supports Dragen and multiple open-source aligners. This enables HiCUP+ users to run Hi-C data processing with several choices of aligners, including HiSAT2 or Dragen which have significant advantages over currently available implementations. Using HiCUP+, the increased accuracy and reduced compute-time in these modern aligners developed for genomic and transcriptomic alignment can now be leveraged in Hi-C analyses. We performed extensive testing and debugging to ensure that these modifications to add compatibility for new aligners do not affect the reproducibility of results compared to the current version of HiCUP when running it with the same parameters.

## Software Implementation

The HiCUP+ pipeline is forked from version 0.8.3 of the original HiCUP pipeline (Wingett et al., 2021) with significant modifications. As with the original pipeline, it is implemented in the Perl and R programming languages. We tested HiCUP+ using Perl version 5.16.3a and R version 3.6.0 on a CentOS 7 Linux system (kernel v 3.10) with samtools version 1.14 and all necessary packages from CPAN and CRAN installed. In accordance with the lesser GNU public license (LGPL-3) that HiCUP is released under, we have released HiCUP+ (https://github.com/hugp-ri/HiCUP-Plus) open-source on the same license and carefully documented the modifications, which can be traced with version control.

The HiCUP+ pipeline is designed to be flexible and backwards compatible for existing HiCUP users. In principle, any HiCUP configuration and input files that runs on HiCUP v0.8.3 will in run with HiCUP+ without modifications and return the same results. Where necessary, the aligner configuration has been checked carefully to make sure that steps necessary to support new aligners do not modify the results of Bowtie or Bowtie2.

When HiSAT2 is called by the mapping subroutine, it uses the same stream-input parameters as Bowtie2, where possible. HiSAT2 is configured to exclude unaligned reads, disable soft-clipping, and avoid multiple alignments. This ensures that, despite some differences to the Bowtie2 command-line interface, the HiSAT2 results are compatible with downstream computations and the only difference is using a newer more accurate mapping algorithm and an indexing scheme with faster performance. This enables using a newer aligner when Dragen is not available.

The main novel feature of HiCUP+ is support for the Illumina Dragen pipeline when a licensed version is available. In particular, it is recommended to use Dragen on the proprietary Illumina Bio-IT platform with the necessary hardware to gain performance improvements using FPGA technology. Dragen is invoked by a shell command via a system call in the Perl script, with clipping parameters disabled for compatible outputs. The logs from the call are passed to stdout with warning messages only displayed in stderr (this differs from Bowtie2 and HiSAT2 which report logs via stderr). The output files were then passed to later steps computed by HiCUP with a custom Perl subroutine to import the summary statistics reported in Dragen log files.

To support Dragen, hard clipping is disabled, reads too short to map are ignored and the reported totals are adjusted, reverse reads are initialized, and pairing uses matches in the read header strings rather than index numbers. This approach maintains compatibility with other aligners while supporting Dragen which does not allow giving index numbers to each read in stream input. Combined, these modifications to the mapping, pairing, and reporting steps ensure full support for Dragen alignment by adding the path to the aligner binary to the PATH environment variable or the configuration file.

## Performance

We have tested the HiCUP+ pipeline on published Hi-C data from the FANTOM6 project (Ramilowski et al., 2020; Agrawal et al., 2021). These datasets were available in raw FASTQ format from public repositories. These were chosen instead of the original Hi-C protocol (Leiberman-Aiden et al., 2009) to demonstrate the HiCUP+ pipeline on large-scale genomics data from modern Illumina platforms.

All jobs were run in parallel on the same dedicated 48-core server separately, ensuring that no other computations were being run. A commercially licensed Dragen node running CentOS 7 with FPGA architecture available was used for all computations, with 48 threads. HiCUP v0.8.3 and HiCUP+ v1.0.1 were both run with alternative aligners, being the only difference in configuration. Input data in raw FASTQ format was downsampled to 1 thousand, 10 thousand, 100 thousand, 1 million, 10 million and 100 million reads respectively and then quality trimmed using the same thresholds (phred score >= Q30 and min-length >= 20) for each run.

As shown in Table 1 for 1 million reads (full results for all samples and file sizes are provided in Supplementary File S1), we clearly demonstrate that aligner runtime and total cpu-hours were substantially lower for HiCUP+ when using HiSAT2 or Dragen compared to the previously supported aligners. It is clear from examining the compute-time of each step that mapping was a rate-limiting step, impacting the performance of the HiCUP pipeline and taking a significant proportion of the total compute-time. Compute-time for truncation and reporting steps in HiCUP+ was not different between alternatives aligners while downstream filtering and deduplication was slower. Therefore, reduced compute-time required by the aligner could be effectively leveraged by the pipeline, with the net compute-time correspondingly improved.

**Table 1.**
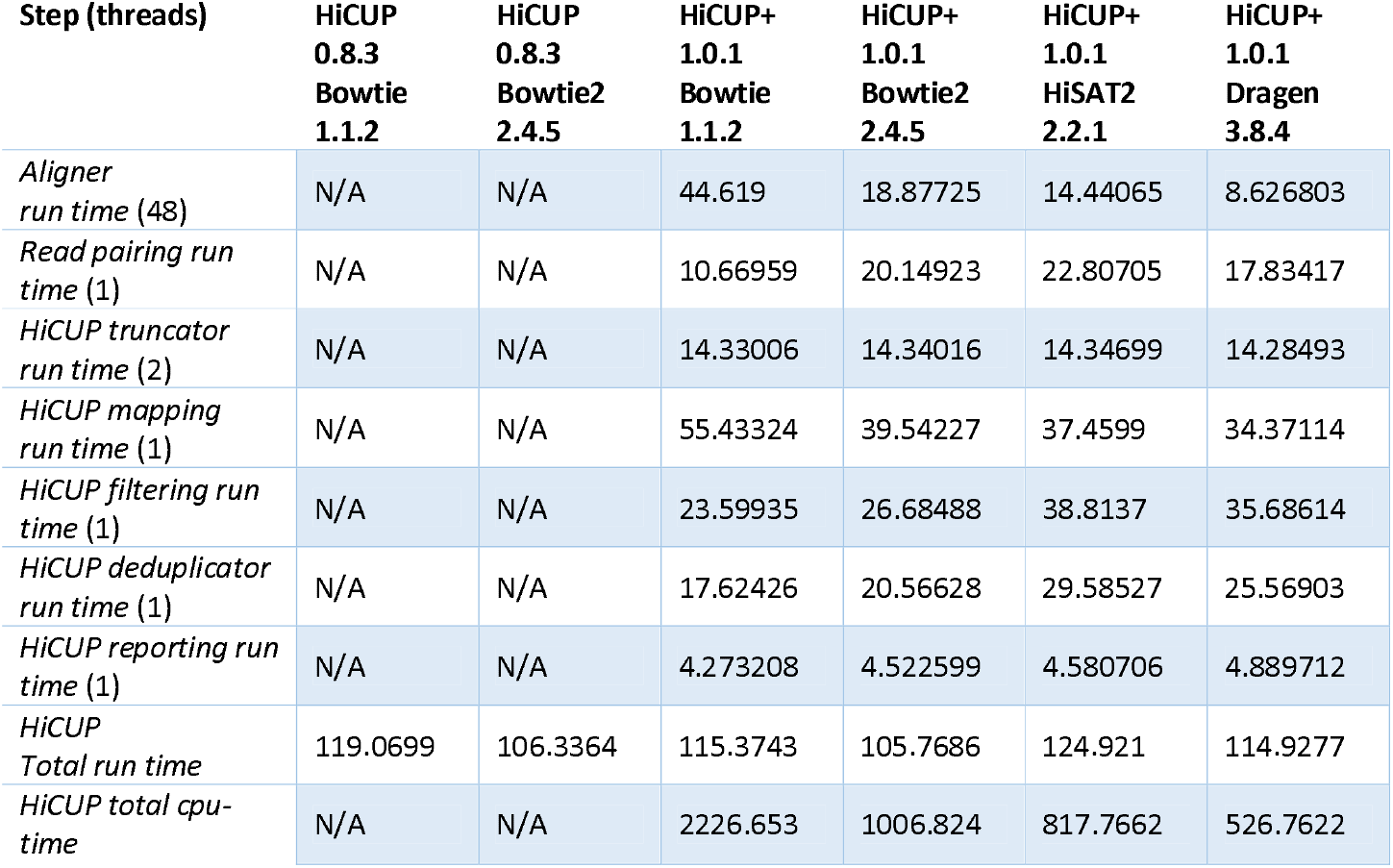
Comparison of compute-time using different aligner. Data shown for input files with 1 million reads using the same 48 core server (time in seconds). Note that compute-time for each step is only reported by HiCUP+ v1.0.1 or later.

| Step (threads) | HiCUP<br>0.8.3<br>Bowtie<br>1.1.2 | HiCUP<br>0.8.3<br>Bowtie2<br>2.4.5 | HiCUP+<br>1.0.1<br>Bowtie<br>1.1.2 | HiCUP+<br>1.0.1<br>Bowtie2<br>2.4.5 | HiCUP+<br>1.0.1<br>HiSAT2<br>2.2.1 | HiCUP+<br>1.0.1<br>Dragen<br>3.8.4 |
| --- | --- | --- | --- | --- | --- | --- |
| *Aligner run time (48)* | N/A | N/A | 44.619 | 18.87725 | 14.44065 | 8.626803 |
| *Read pairing run time (1)* | N/A | N/A | 10.66959 | 20.14923 | 22.80705 | 17.83417 |
| *HiCUP truncator run time (2)* | N/A | N/A | 14.33006 | 14.34016 | 14.34699 | 14.28493 |
| *HiCUP mapping run time (1)* | N/A | N/A | 55.43324 | 39.54227 | 37.4599 | 34.37114 |
| *HiCUP filtering run time (1)* | N/A | N/A | 23.59935 | 26.68488 | 38.8137 | 35.68614 |
| *HiCUP deduplicator run time (1)* | N/A | N/A | 17.62426 | 20.56628 | 29.58527 | 25.56903 |
| *HiCUP reporting run time (1)* | N/A | N/A | 4.273208 | 4.522599 | 4.580706 | 4.889712 |
| *HiCUP Total run time* | 119.0699 | 106.3364 | 115.3743 | 105.7686 | 124.921 | 114.9277 |
| *HiCUP total cpu-time* | N/A | N/A | 2226.653 | 1006.824 | 817.7662 | 526.7622 |

Furthermore, changes to the HiCUP+ pipeline did not impact performance of existing aligners. Performance of HiCUP+ v1.0.1 using no different to release v0.8.3 of HiCUP when using Bowtie *(t* = 1.0352, *df* = 3.4895, *p* = 0.3669) or Bowtie2 (*t* = -0.54659, *df* = 3.3346, *p* = 0.6191) respectively, as shown in Table 2 for 1 million reads in 3 samples. Thus it is possible to support new aligners without negatively affecting those already supported.

**Table 2.** Performance of HiCUP and HiCUP+ are very similar for supported aligners. Bowtie and Bowtie2 had near identical compute-time when running HiCUP v0.8.3 or HiCUP+ 1.0.1 with 1 million reads input for 3 samples.

| Sample | HiCUP<br>0.8.3<br>Bowtie<br>1.1.2 | HiCUP<br>0.8.3<br>Bowtie2<br>2.4.5 | HiCUP+<br>1.0.1<br>Bowtie<br>1.1.2 | HiCUP+<br>1.0.1<br>Bowtie2<br>2.4.5 | HiCUP+<br>1.0.1<br>HiSAT2<br>2.2.1 | HiCUP+<br>1.0.1<br>Dragen<br>3.8.4 |
| --- | --- | --- | --- | --- | --- | --- |
| *Sample DRR179523* | 119.0699 | 106.3364 | 115.3743 | 105.7686 | 124.921 | 114.9277 |
| *Sample DRR179524* | 115.6033 | 103.5923 | 117.4667 | 106.1857 | 121.2312 | 111.8521 |
| *Sample DRR179525* | 117.4641 | 107.8834 | 115.555 | 108.2768 | 125.6476 | 111.7277 |

The total compute time for HiCUP and HiCUP+ were compared across different input file sizes for 3 replicate samples. As shown in Figure 1, the Dragen and HiSAT2 aligners had reasonable performance on large files. While they have poorer performance on small files due to longer overhead times, they are optimised for large FASTQ input files as typically required by genomic analysis. In particular, the Dragen aligner significantly reduces run-time requirements for the pipeline on large input files. HiSAT2 had similar total runtime to Bowtie2 allowing refinements in the newer algorithm to be used without much additional runtime. As previously observed (Table 2), the runtime for HiCUP v0.8.3 was very similar for supported aligners (Bowtie and Bowtie2).

**Figure 1.**
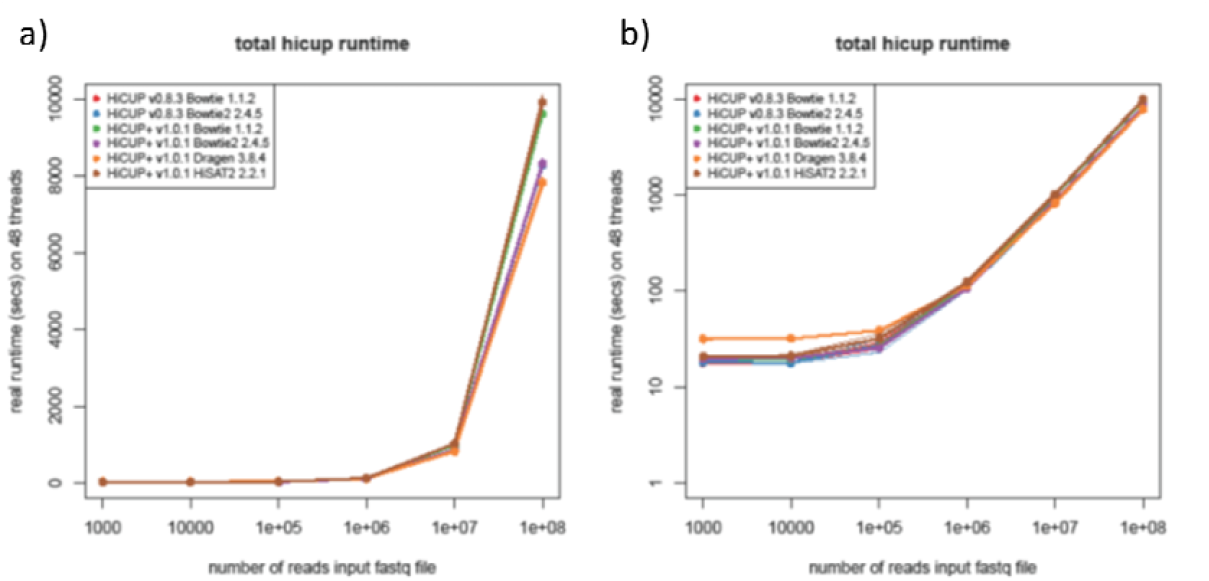
Choice in aligner affects total runtime. Run time for HiCUP and HiCUP+ with different aligners. Total runtime for each pipeline and aligner combination on linear (right) and log-scale (left) are shown for various input file sizes. Results were reported as a mean of 3 samples with error margins of 1 standard deviation. Results for HiCUP and HiCUP+ were near identical when running Bowtie or Bowtie2 for each aligner respectively. Dragen and HiSAT2 had longer runtimes on smaller files due to overhead set up time which translated into faster runtime on larger files.

Examining the runtime for each step in the HiCUP+ pipeline (Figure 2), the performance on larger files and difference between aligners can be observed (raw data is provided in Supplementary File S1). The truncation pre-processing step (Figure 2a) depends only on input file size and was no different between aligners, as expected. Runtime for mapping (Figure 2b) was longer for Dragen and HiSAT2 with smaller input files due to longer set up time but this is offset by improved performance with larger files. The post-processing steps of filtering (Figure 2c) and deduplication (Figure 2d) were not very different between aligners. Marginally longer runtime for Dragen and HiSAT2 partially offsets the performance benefits of faster mapping but can be explained by larger output files and higher mapping rate as shown below as these steps depend on the input file size (output from mapping). Note that linear trends on a log-log scale for larger files suggest that performance follows a power-law distribution. The reporting step (Figure 2e) to generate Rmarkdown reports from summary statistics had a constant runtime irrespective of raw input files or aligner chosen.

**Figure 2.**
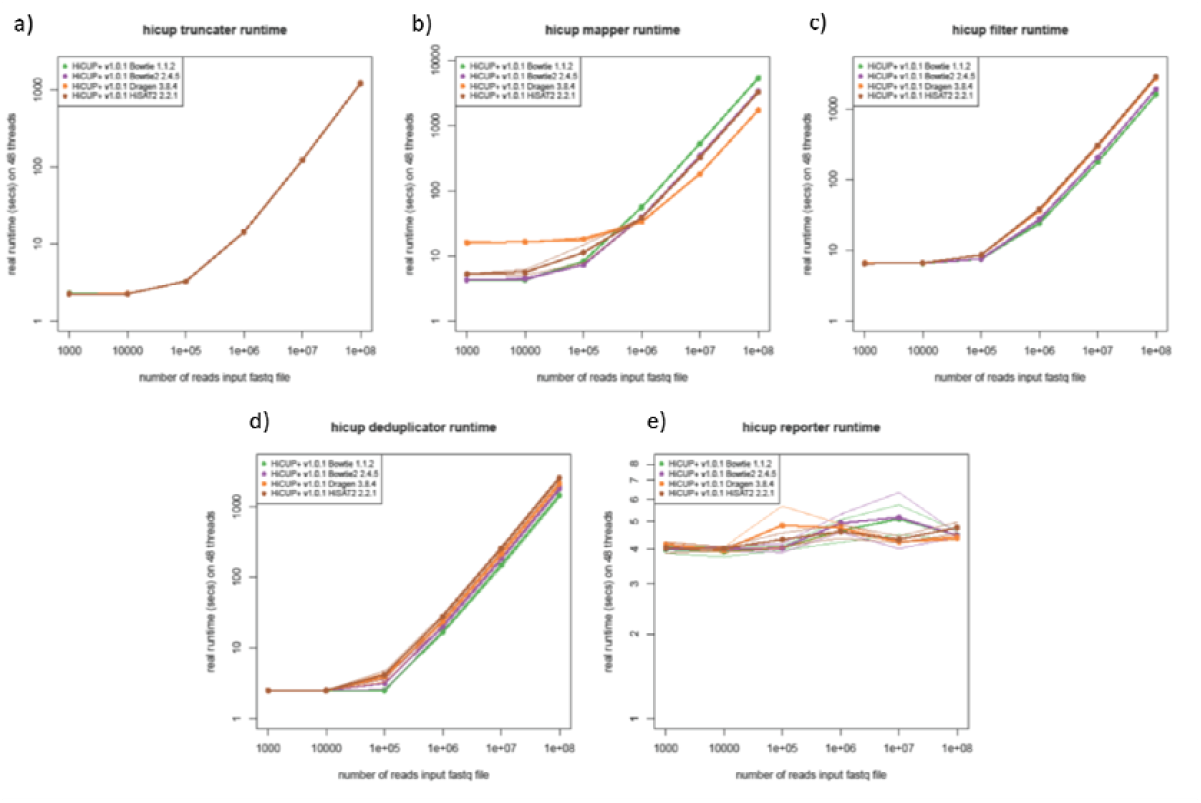
Changing aligners affects performance of downstream steps with larger files. HiCUP+ was run with 4 aligners with runtime reported for each step of the pipeline at various input file sizes as a mean of 3 samples with error margins of 1 standard deviation.

Note that several steps of the pipeline are multi-threaded to reduce the real runtime of the pipeline. Specifically, truncation is dual-threaded, running 2 processes on each read1 and read2 input file concurrently. Threads for mapping in parallel can be configured and 48 threads were used for all aligners. Adjusting for multithreading the compute time (CPU-time) for the HiCUP+ pipeline and all steps can be calculated (Figure 3). Each step exhibits the same trends as observed above but accounting for multi-threaded processes, it is clear that the total compute-time (Figure 3a) is limited by mapping compute time (Figure 3c) and can be specifically attributed to the alignment step (Figure 4).

**Figure 3.**
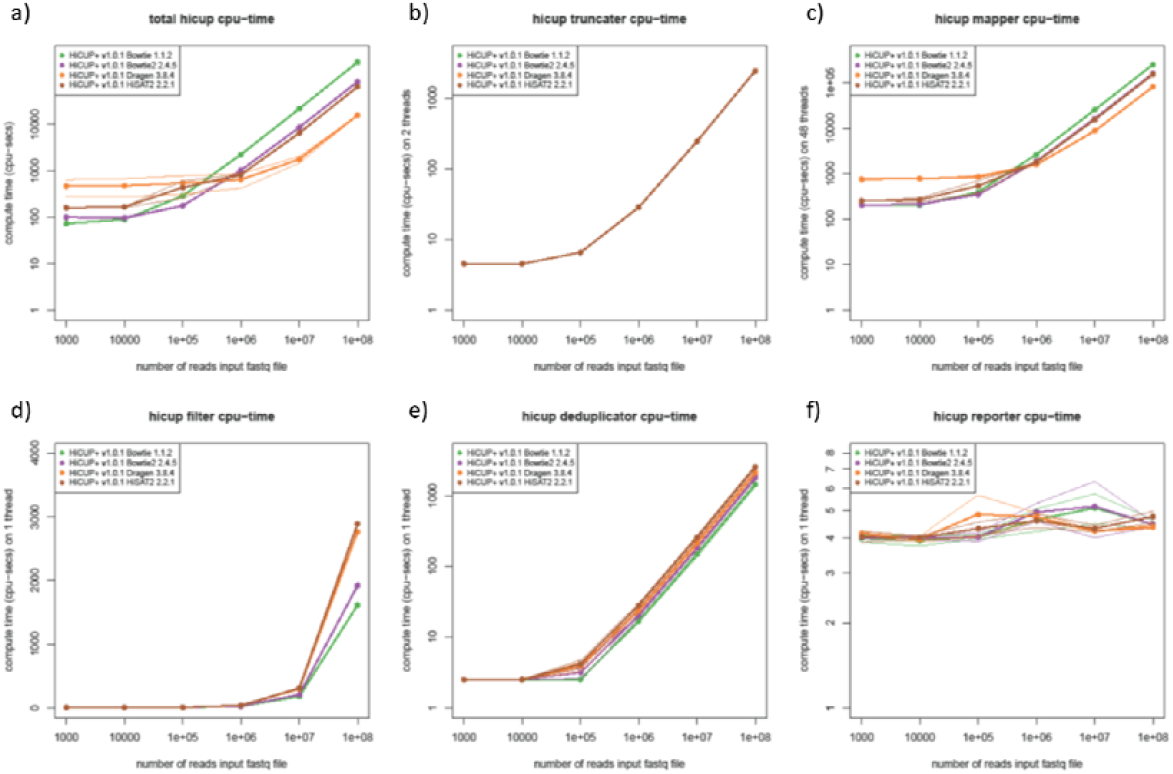
Compute-time for HiCUP+ is rate-limited by mapping. All four aligners supported by HiCUP+ were run with each step of the pipeline timed. Compute-time of each step is shown for various input file sizes, accounting for multi-threaded steps. Results were reported as a mean of 3 samples with error margins of 1 standard deviation. Trends in total HiCUP+ runtime can be attributed to the mapping step as it is the most computationally resource intensive.

**Figure 4.**
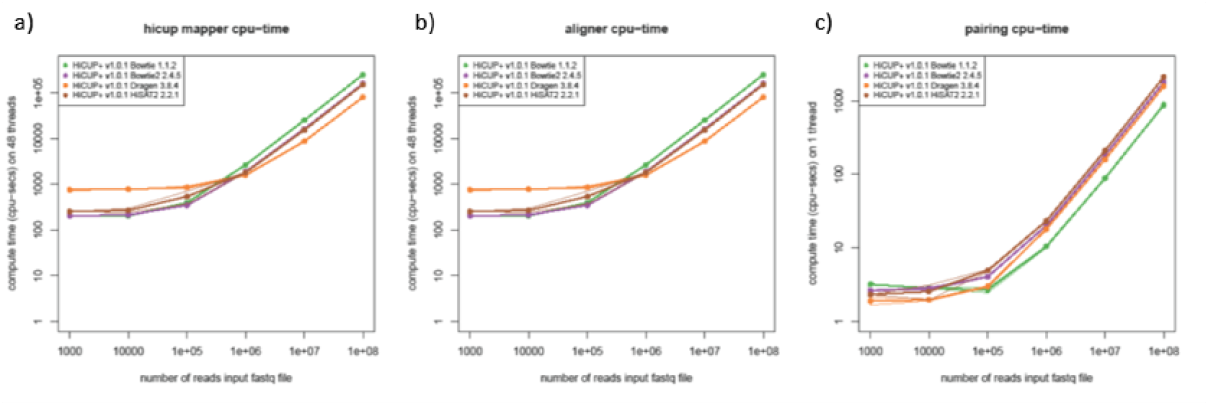
Adopting alternative aligners yields performance improvements in the alignment subroutine. All four aligners supported by HiCUP+ were run with the aligner call of the pipeline timed. Compute-time of the aligner call compared to the other main step (pairing) in the HiCUP+ mapper is shown for various input file sizes, accounting for multi-threaded steps. Results were reported as a mean of 3 samples with error margins of 1 standard deviation. This demonstrates that HiSAT2 and Dragen are optimised for large files and adopting either of these as an alternative aligner improves performance in this crucial step.

As mapping is the most computationally demanding step, it is the logical choice for optimisation. Adopting Dragen or HiSAT2, reduces the compute-time for alignment (Figures 4b and 4e) with HiSAT2 having marginally faster performance than Bowtie2 and Dragen drastically reducing compute-time for alignment. These benefits improve the performance of the mapping step and total pipeline in turn. While real runtime can be reduced by computing alignments in parallel, these require large remote computing resources. Furthermore resource-intensive alignment algorithms are costly on cloud-computing services for large input files (as shown in Table 3). Larger mapped outputs from these aligners also increases compute-time of pairing (Figures 4c and 4f) but this step is less resource-intensive and must be performed single-threaded to ensure correct order of reads. This demonstrates the benefit of adopting new aligners by addressing most computationally demanding step of the pipeline HiSAT2 and Dragen are both faster than existing options for `hicup_mapper`.

**Table 3.**
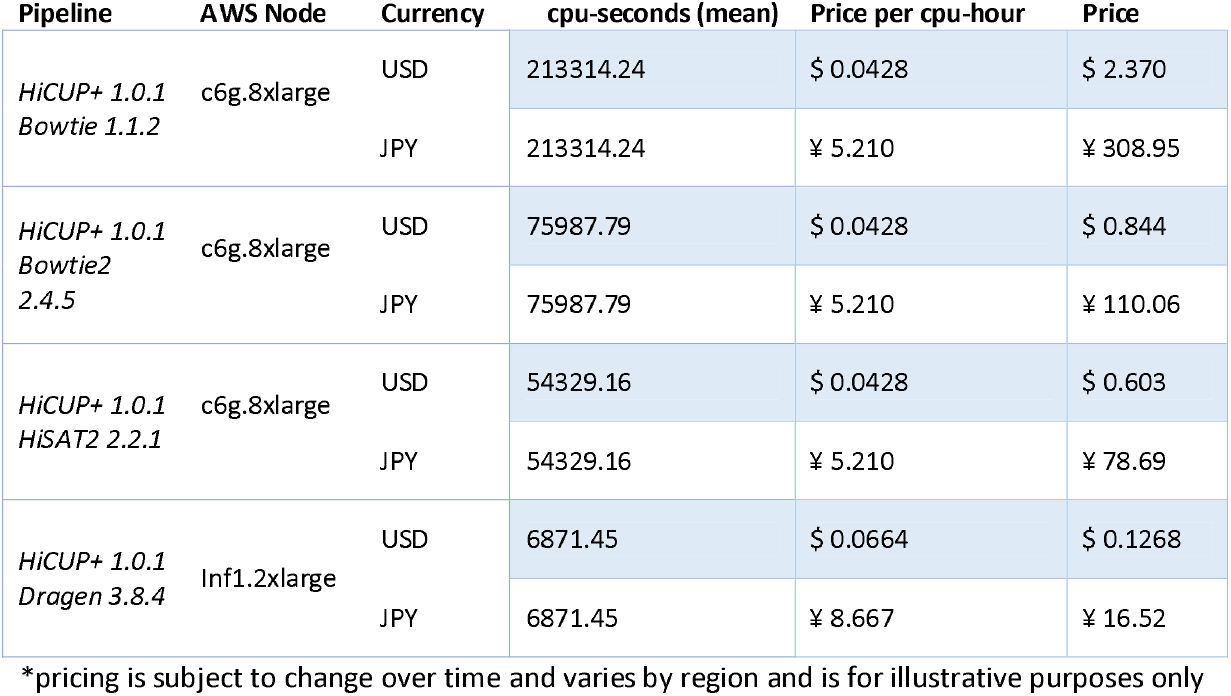
Costing on cloud compute services (prices from AWS EC2 instances for running 100 million reads, in the Asia Pacific Northeast region with exchange rates at the time of writing).

The `hicup_mapper` subroutine has 2 time-consuming operations: first it aligns each read to the genome independently and then after unmapped reads are discarded, the reads are paired (Wingett et al., 2015). A notable change in HiCUP+ is that the pairing step after alignment with

Dragen uses matching FASTQ file headers, rather than index numbers assigned during alignment for other aligners using stream inputs. In order to do this efficiently, the sorted output in Dragen must be disabled to ensure the aligned reads are in the same order as the FASTQ files. In figure 4, we demonstrate that the alignment step specifically is improved by adopting HiSAT2 or especially the Dragen aligner. Since other operations have similar performance for each aligner, we demonstrate that reducing mapping runtime improves the performance when running the entire HiCUP+ pipeline. The index numbers are retained for full backwards compatibility for other aligners, although similar runtime for pairing suggests no impacts on performance from changing to matching strings. As shown below, longer runtime for pairing with HiSAT2 and Dragen can be explained by larger outputs of uniquely mapped reads due to higher mapping accuracy.

As shown in Figure 5, the overall compute-time for HiCUP+ is significantly reduced when using Dragen alignment and this can be largely attributed to the alignment step (shown in Figure 4). Here the `hicup_mapper` is plotted separately for the aligner call and pairing steps. The compute-time for truncation and mapping increases for larger files, while the downstream processing of the aligner output is a fixed time regardless of input file size. Therefore, for larger files in particular, the majority of compute-time is consumed by running alignment when using the Bowtie aligner (shown in Fig 5d). Newer aligners Bowtie2, HiSAT2, or Dragen each further reduce the compute-time of the alignment step and in turn reduce the proportion of compute-time required for mapping within the pipeline. The improvements in larger files are therefore worth the increased overhead compute time required to set up alignment as shown for smaller input files. This is likely because these aligners are optimised to set up reference indexing systems and parallel computing to increase performance on large inputs. Therefore it is recommended to use newer aligners of HiSAT2 or Dragen when available to run the pipeline more efficiently with fewer computational resources and shorter wait times.

**Figure 5.**
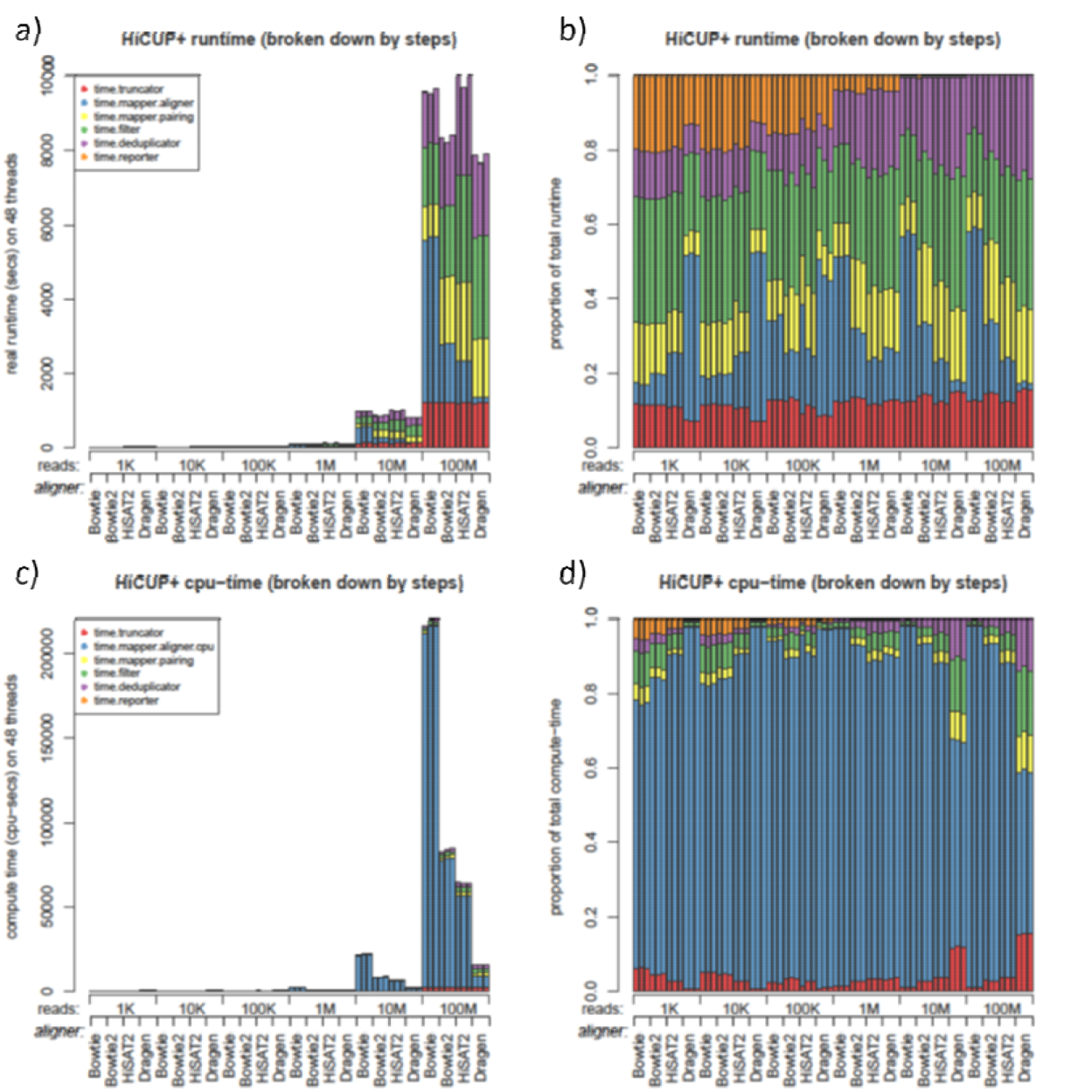
Aligner choice enables addressing computational performance of the rate-limiting mapping step. All four aligners supported by HiCUP+ were run with each step of the pipeline timed. The mapping step is divided into alignment and pairing subroutines. Compute-time of each step is shown for various input file sizes, accounting for multi-threaded steps. Results were reported for the same 3 replicate samples for each aligner. Total runtime is shown for each step in absolute time (a) and proportions (b) of the total for each run. Similarly compute-time accounting for multi-threaded steps is shown in compute-seconds (c) across all CPUs used and proportions (d) of total compute-time. For larger files, the alignment step with Bowtie or Bowtie2 takes up the vast majority of compute-time. Realtime turnaround and computational efficiency can thus be improved by using HiSAT2 or Dragen which reduces the compute-time required for this step.

## Accuracy and Reproducibility

It is notable that these improvements in performance, with reduced compute-time and memory requirements, do not come at the expense of accuracy. In fact the opposite occurs, with newer aligners having refined mapping accuracy. This impacts performance of downstream post-processing, owing to the larger outputs from higher accuracy and mapping rates.

The HiSAT2 (Kim et al., 2019) aligner has been previously shown to be highly accurate with improvements over Bowtie2 (Musich et al., 2021). Dragen (Illumina, Inc., 2021) has been further optimised for accurate variant calling in difficult to map regions of the genome and improved over previous versions in the PrecisionFDA Truth challenge V2 (Illumina, Inc., 2020a; Illumina, Inc., 2020b; Olson et al., 2021; Wagner et al., 2021). Introducing these aligners to the HiCUP+ pipeline allows these highly accurate mapping tools to be used in existing Hi-C data processing workflows. We anticipate that the improvements in handling difficult to map regions and improving mapping accuracy will in turn translate to improved results in Hi-C analysis.

Furthermore, the modifications to HiCUP+ do not alter the reproducibility of existing workflows using the Bowtie or Bowtie 2 aligners. We confirmed that the `hicup_mapper` output for HiCUP v0.8.3 and HiCUP+ v1.0.1 was identical with respect to the mapping and pairing rates (as shown in Table 4). These aligners are retained to support legacy configurations and can run on systems where only these aligners are installed. We also demonstrate that the changes to filtering and reporting steps to accommodate new aligners does not change the results when using an aligner that was already supported (as shown in Table 5). The number of valid pairs and di-tags in the output results were the same using either Bowtie or Bowtie2 with both HiCUP v0.8.3 and HiCUP+ v1.0.1. Thus we have ensured that existing functionality was completely unaffected by alternations to the HiCUP source code and the pipeline is fully-compatible with the original aligners.

**Table 4.**
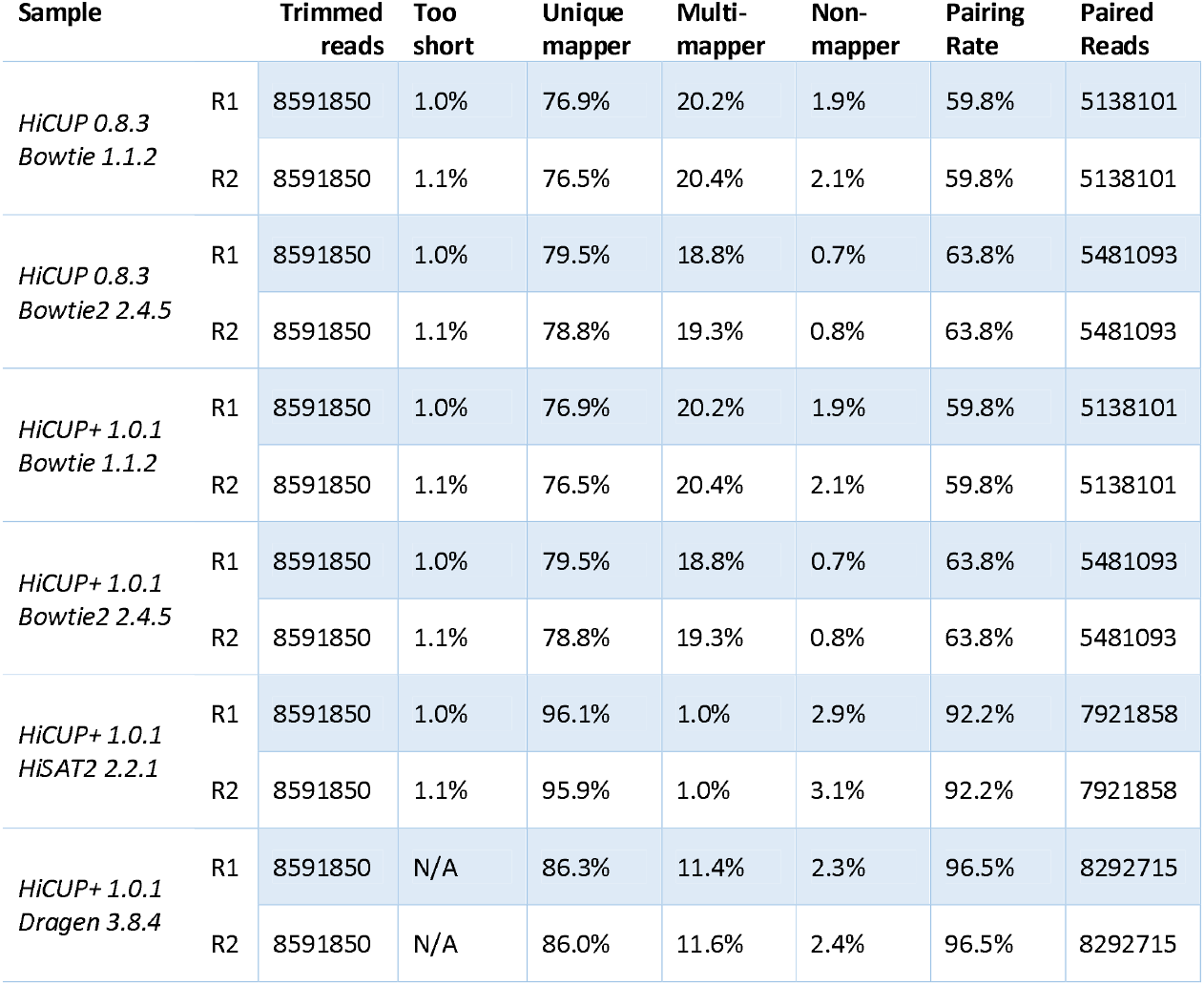
Results from HiCUP and HiCUP+ mapping summary statistics for supported aligners. Bowtie and Bowtie2 had exactly the same results when running HiCUP v0.8.3 or HiCUP+ 1.0.1 with 10 million raw reads input for 3 samples.

**Table 5.** Results from HiCUP and HiCUP+ reporting summary for supported aligners. Bowtie and Bowtie2 had exactly the same results when running HiCUP v0.8.3 or HiCUP+ 1.0.1 with 10 million raw reads input for 3 samples.

| Sample |  | Truncated reads | Length remove (mean) | Mapped rate | Valid Pairs | Unique Pairs | Unique Trans | Di-tags passing filter |
| --- | --- | --- | --- | --- | --- | --- | --- | --- |
| *HiCUP 0.8.3*<br>*Bowtie 1.1.2* | R1 | 726199 | 30.76 | 59.80% | 72.01% | 99.96% | 66.10% | 43.05% |
|  | R2 | 721230 | 30.53 | 59.80% | 72.01% | 99.96% | 66.10% | 43.05% |
| *HiCUP 0.8.3*<br>*Bowtie2 2.4.5* | R1 | 726199 | 30.76 | 63.79% | 71.98% | 99.96% | 66.12% | 45.90% |
|  | R2 | 721230 | 30.53 | 63.79% | 71.98% | 99.96% | 66.12% | 45.90% |
| *HiCUP+ 1.0.1*<br>*Bowtie 1.1.2* | R1 | 726199 | 30.76 | 59.80% | 72.01% | 99.96% | 66.10% | 43.05% |
|  | R2 | 721230 | 30.53 | 59.80% | 72.01% | 99.96% | 66.10% | 43.05% |
| *HiCUP+ 1.0.1*<br>*Bowtie2 2.4.5* | R1 | 726199 | 30.76 | 63.79% | 71.98% | 99.96% | 66.12% | 45.90% |
|  | R2 | 721230 | 30.53 | 63.79% | 71.98% | 99.96% | 66.12% | 45.90% |
| *HiCUP+ 1.0.1*<br>*HiSAT2 2.2.1* | R1 | 726199 | 30.76 | 96.52% | 60.95% | 99.99% | 81.23% | 58.82% |
|  | R2 | 721230 | 30.53 | 96.52% | 60.95% | 99.99% | 81.23% | 58.82% |
| *HiCUP+ 1.0.1*<br>*Dragen 3.8.4* | R1 | 726199 | 30.76 | 92.20% | 68.35% | 99.96% | 70.51% | 63.00% |
|  | R2 | 721230 | 30.53 | 92.20% | 68.35% | 99.96% | 70.51% | 63.00% |

These results demonstrate the benefits of adopting a newer more accurate alignment algorithm. As shown in Table 4, HiCUP+ returned a higher mapping and pairing rate for the same sample when using HiSAT2 or Dragen than existing aligners for `hicup_mapper` (see Supplementary File S2 for full output). Reads too short to be mapped are not reported for Dragen as this is only enabled for aligners that support stream inputs. The uniquely mapping rates were higher at 95.9% and 86.0% for HiSAT2 and Dragen respectively. Similarly, the pairing rates were higher at 92.2% and 95.5% for HiSAT2 and Dragen respectively.

Both alternative aligners returned more paired reads in the output BAM file from `hicup_mapper` than Bowtie or Bowtie2. These larger outputs (approximately 40-60% more paired reads) account for the differences in performance in post-processing discussed above. As shown in Table 5, these higher rates of paired reads generated higher number of di-tags passing all post-processing filters with 58.8% and 63.0% for HiSAT2 and Dragen respectively (see Supplementary File S3 for full output). This is because HiCUP+ requires uniquely mapped reads with a high confidence. Therefore we demonstrate how adoption of newer more accurate alignment algorithms is beneficial for generating high-resolution Hi-C datasets and explain the impacts of these aligners on performance with longer compute-time for downstream post-processing scripts.

## Significance

This combined pipeline has several advantages over currently available alternatives. Compared to other sequence processing pipelines, using HiCUP accounts for biases in restriction enzyme digests with a custom reference. Alignment is a rate-limiting step in the pipeline, requiring multi-threaded computations over a significant period of time on large computational resources. By integrating modern aligners into this pipeline, we were able to increase the flexibility of HiCUP and leverage the performance and accuracy improvements provided by Dragen or HiSAT2.

The Dragen aligner is particularly advantageous because this allows taking advantage of significant performance improvements on the commercially available Bio-IT platform (Illumina Inc., 2021) with hardware acceleration using field-programmable gate array technology (FPGA). The licensed Dragen pipeline also provides rich summary information, including built-in quality checking (QC) tools and summary statistics. The HiCUP+ pipeline enables simultaneously generating this summary information along with the HiCUP post-processing and summary computations tailored for Hi-C data. Both are supported by HiCUP and MultiQC modules (Ewels et al., 2015).

Several alternative aligners are supported, including Bowtie2 and HiSAT2, which do not require commercial licensing of Dragen or custom hardware. These can be configured with backwards compatibility with the original HiCUP release. Any configuration file written for the previous version of HiCUP v0.8.3 will also be compatible with HiCUP+ and return exactly the same results if the same aligner is specified. As with the original pipeline, the aligner can be specified in the configuration pipeline or aligners in the PATH environment variable will be used if available, defaulting to Dragen, HiSAT2, or Bowtie2 in order of priority. It is therefore straightforward to migrate existing pipelines using the popular HiCUP pipeline to use HiCUP+ by installing the new aligner and adding it to the configuration.

By supporting newer alternative aligners, with faster performance and higher mapping accuracy, the HiCUP+ pipeline enables high quality Hi-C data processing with reduced computational resources. This has several inherent advantages in sequencing applications, particularly with large-scale sequencing data. Higher mapping accuracy not only ensures that each read is mapped correctly to the right locus but also loses fewer unaligned or multi-mapped reads. HiSAT2 and Dragen are able to accurately align reads with poorer quality sequence and more mismatches to the reference. In particular, these aligners were designed to support splice-variants in RNA sequence data and structural variants in genomic sequence data.

These aligners also perform significantly faster computations than Bowtie2, with overall compute-hours reduced by adopting one of these aligners. Reducing cpu-time allows running them on large-scale datasets and reducing costs to run pipelines in cloud computing environments. As they can be run in parallel using multiple threads, which further reduces the real run-time, this allows for rapid turn-around time. These performance improvements are advantageous to handle a large volume of samples and report results quickly, particularly for commercial sequencing services.

Together, we believe these improved features are clearly beneficial to the wider research community. HiCUP+ has been publicly released on an LGPL-3 license which permits reuse, modification, and commercial use with attribution to the original authors of HiCUP (Wingett et al., 2015). The additional aligners available gives researchers the flexibility to choose from multiple alternatives for processing Hi-C data to best address the specific needs of their project. Crucially it is possible to introduce improvements in genomic DNA mapping technologies to Hi-C analyses without sacrificing computational speed or quality of results.

## Concluding Remarks

In summary, we have integrated the popular HiSAT2 and Dragen aligners into the HiCUP pipeline, allowing large scale Hi-C datasets to be efficiently and accurately mapped and processed in a reproducible fashion. With respect to the original developers, we have developed this custom pipeline using version control and released it open-source on the same license, with the new optional configurations fully-documented. This allows greater flexibility in the HiCUP pipeline, with a greater range of aligners for users to choose from depending on availability and the needs of the project. Integrating Dragen specifically has significant advantages because it has improved performance. Both HiSAT2 and Dragen have higher mapping accuracy than previously available aligners which yields higher resolution processed Hi-C datasets. We have ensured that the pipeline can reproduce results from the previous version using previously supported aligners. This ensures that only the necessary changes to support new aligners have been made without altering the results of other steps when adding compatibility to Dragen output formats. To benefit the research community, we have released the open-source HiCUP+ pipeline with improved Hi-C data processing capabilities for fast and accurate result.

## Acknowledgements

We wish to acknowledge Steven Wingett (The Babraham Institute, Cambridge, UK) and colleagues who developed the original HiCUP pipeline on an open-source license. We thank Kai Battenberg (RIKEN CSRS, Yokohama, Japan) for advice on parsing specific file formats in Perl. Saumya Agrawal (RIKEN IMS, Yokohama, Japan), Yuki Kawasaki (H.U. Group Research Institute, Akiruno, Japan), and Kenichiro Kori (H.U. Group Research Institute/SRL Inc., Akiruno, Japan) gave helpful advice on the Hi-C experimental technique. Ayumi Kuga (H.U. Group Research Institute/SRL Inc., Akiruno, Japan) gave advice on cloud computing costs.

## Code availability

The HiCUP+ pipeline is publicly released on an LGPL-3 license (which allows modification and commercial use on certain conditions). The source code is available at the following URL:

https://github.com/hugp-ri/HiCUP-Plus

The version 1.0.1 release discussed in this manuscript is archived: https://github.com/hugp-ri/hicup-plus/releases/tag/v1.0.1

Updated documentation on this software is available at this webpage:

https://hicup-plus.readthedocs.io/en/latest/

### Data availability

All data used has been previously published (Ramilowski et al., 2020; Agrawal et al., 2021) and is available in publicly accessible repositories:

https://fantom.gsc.riken.jp/6/datafiles/Hi-C_public_repository/

https://ddbj.nig.ac.jp/resource/bioproject/PRJDB7993

## Supplementary Figures

**Figure S1.**
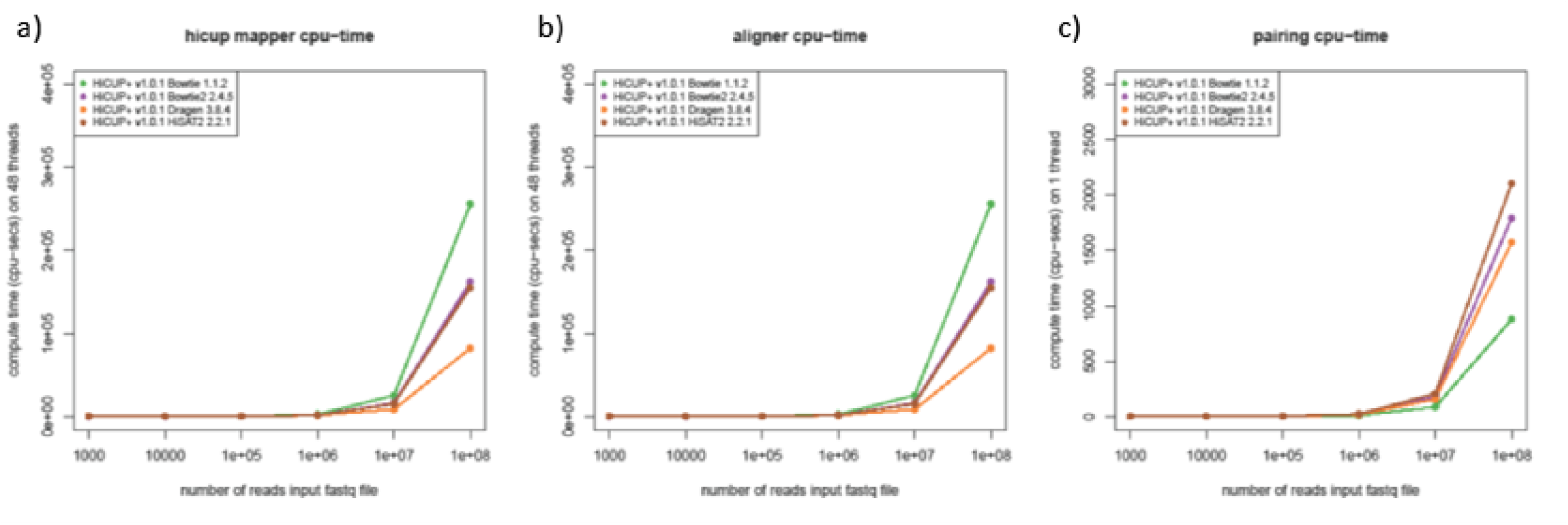
Adopting alternative aligners yields performance improvements in the alignment subroutine. All four aligners supported by HiCUP+ were run with the aligner call of the pipeline timed. Compute-time of the aligner call compared to the other main step (pairing) in the HiCUP+ mapper is shown for various input file sizes, accounting for multi-threaded steps. Results were reported as a mean of 3 samples with error margins of 1 standard deviation. All y-axes are shown in linear scale (log-scale is shown in Fig. 4).

